# EMBED: Essential Microbiome Dynamics, a dimensionality reduction approach for longitudinal microbiome studies

**DOI:** 10.1101/2021.03.18.436036

**Authors:** Mayar Shahin, Brian Ji, Purushottam D. Dixit

**Affiliations:** Department of Physics, University of Florida, Gainesville, FL 32611; Physician-Scientist Training Pathway, Department of Medicine, UCSD, San Diego, CA 92103; Genetics Institute, University of Florida, Gainesville, FL 32611; Department of Chemical Engineering, University of Florida, Gainesville, FL 32611

## Abstract

Dimensionality reduction can offer unique insights into high dimensional microbiome dynamics by leveraging collective abundance fluctuations of multiple bacteria driven by similar ecological perturbations. However, methods providing lower-dimensional representations of microbiome dynamics both at the community and individual taxa level are not currently available. To that end, we present EMBED: **E**ssential **M**icro**B**iom**E D**ynamics, a probabilistic non-linear tensor factorization approach. Similar to normal mode analysis in structural biophysics, EMBED infers ecological normal modes (ECNs), which represent the unique orthogonal modes capturing the collective behavior of microbial communities. A very small number of ECNs can accurately approximate microbiome dynamics across multiple data sets. Inferred ECNs reflect specific ecological behaviors, providing natural templates along which the dynamics of individual bacteria may be partitioned. Moreover, the multi-subject treatment in EMBED systematically identifies subject-specific and universal abundance dynamics that are not detected by traditional approaches. Collectively, these results highlight the utility of EMBED as a versatile dimensionality reduction tool for studies of microbiome dynamics.

## Introduction

Advances in sequencing have enabled the characterization of host-associated microbiomes at unprecedented resolution^1,2^. In contrast to static cross-sectional snapshots of these ecosystems, longitudinal studies offer unique insights into the biological processes structuring microbial ecosystems within individual hosts. For example, recent longitudinal studies on gut microbiome have elucidated the determinants of microbiome colonization in early childhood^3,4^, the effects of the microbiome on outcomes following bone-marrow transplant^5^, and the recolonization of microbial communities following antibiotic perturbation^6–11^.

Yet, understanding how the microbiome changes in response to environmental perturbations such as host diet variation^12,13^ and antibiotic administration^10,11^ remains challenging. This is because of the enormous organizational complexity of these ecosystems, comprising thousands of individual bacterial taxa whose abundances vary substantially across space and time^12,14–17^ and across biological replicates^18^. Additionally, technical sequencing noise can seriously confound true abundance changes^15,19,20^. For example, technical noise is likely to be the most dominant factor in the observed abundance variability in more than half the bacterial taxa in longitudinal gut microbiome studies^15^ and likely remains a significant contributor for all measured taxa.

Despite this complexity, recent work suggests that abundances of individual bacterial species fluctuate with collective responses to perturbations^10–13^. Therefore, the high dimensional dynamics of the microbiome could potentially be understood as a dynamics of a few collective variables on a manifold of a much smaller dimension^21^. Indeed, approaches such as multidimensional scaling that embed microbiome samples on a smaller dimensional manifold are popular^22–24^. However, these methods only identify shifts at the community level^18^. Crucially, these methods do not account for temporal correlations in abundances of individual bacterial taxa and variability across subjects. Other approaches based on low rank factorization of log-transformed data^18,25^ do not account for technical measurement noise and the longitudinal nature of the data. Therefore, they may not distinguish between true abundance changes and sampling noise, potentially leading to inconsistent approximations of microbiome compositions and mixed temporal modes.

To address these pressing issues, we present EMBED: **E**ssential **M**icro**B**iom**E D**ynamics. EMBED is a probabilistic non-linear tensor factorization-based dimensionality reduction method specifically tailored to model longitudinal microbiome data. EMBED identifies a set of unique and orthogonal bases which we call *Ecological Normal Modes* (ECNs) and taxa- and subject-specific loadings that quantify the importance of individual ECNs on the abundance of taxa across subjects. EMBED has several salient features. First, EMBED utilizes the exponential Gibbs-Boltzmann distribution, which captures the extensive variability of the species abundances within microbiomes^17^. Second, by restricting the number of ECNS to be low, EMBED can provide a low-dimensional description of the community by filtering out unimportant fluctuations in the data. Third, ECNs are inferred using a probabilistic model that accounts for sequencing noise inherent in all microbiome studies^15^. Fourth, similar to the normal modes in structural biology^26^, ECNs represent statistically independent modes of collective abundance changes. Fifth, the explicit multi-subject treatment in EMBED systematically identifies universal and subject-specific dynamical behaviors and bacterial taxa that exhibit that behavior.

Using synthetic data and several publicly available longitudinal data sets^10–14^, we show that EMBED-based low-dimensional approximation of microbial community dynamics is accurate and robust to sequencing noise, underscoring the low-dimensional nature of microbiome dynamics. Using synthetic data, we show that EMBED infers statistically independent dynamical modes. Using two datasets that encompass major ecological perturbations including dietary changes^13^, and antibiotic administration^10^, we show that the identified ECNs reflected specific ecological behaviors and serve as templates to reconstruct the dynamics of individual bacterial taxa. The loadings identify universal and subject-specific bacterial taxa dynamics. These results show that EMBED will be an important dimensionality reduction tool to decipher collective dynamical behaviors within the microbiome.

## Results

### EMBED identifies reduced-dimensional descriptors for longitudinal microbiome dynamics

In EMBED (Fig. 1A and **Supplementary Information**), we model microbial abundance counts *n*_*os*_(*t*) (Operational taxonomic unit, OTU “*o*”, subjects “*s*”, and time points “*t*”) as arising from a multinomial distribution. The likelihood of observing the data is given by:

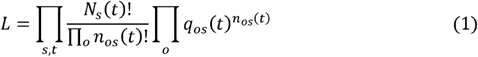

where *N*_*s*_ (*t*) = Σ_*o*_ *n*_*os*_ (t) is the total read count on a given day *t* for subject *s*. The probabilities *q*_*os*_ (t) are modeled as a Gibbs-Boltzmann distribution^27^

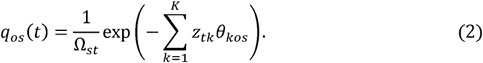

**Figure 1.**
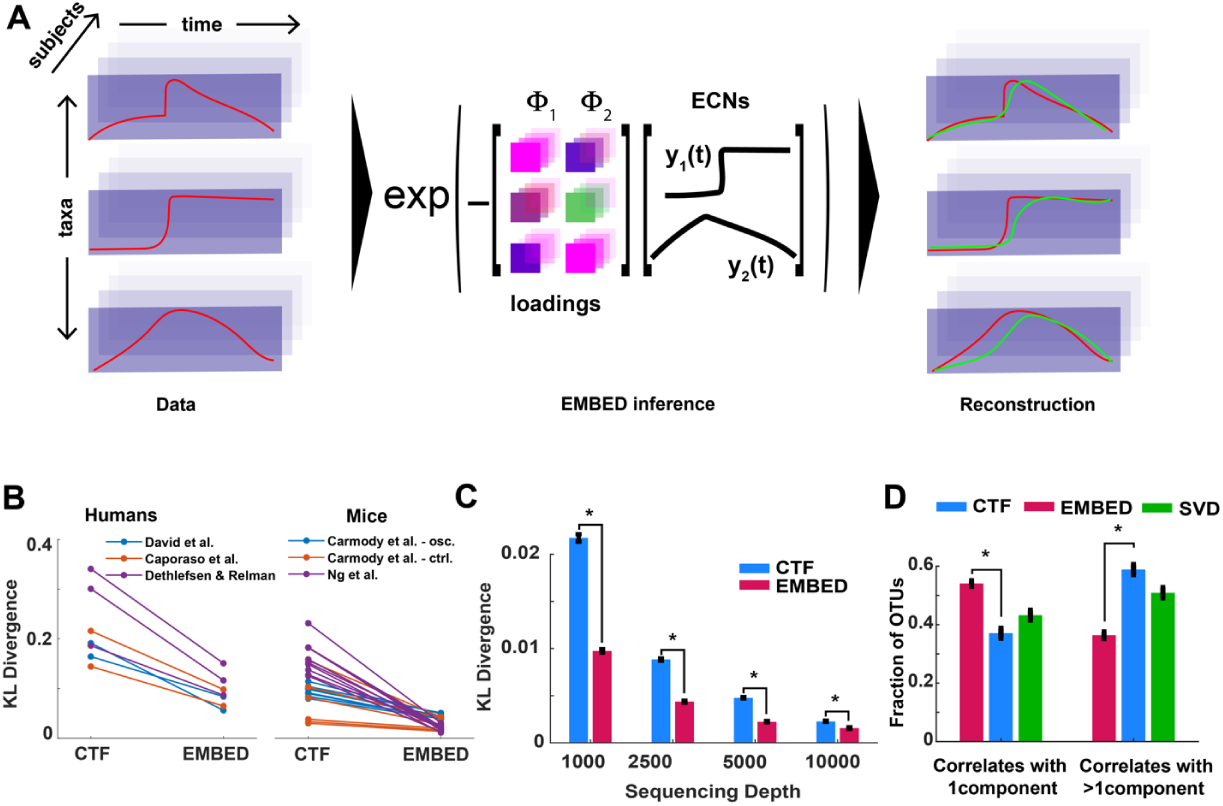
(A) Schematic of the EMBED approach. In the schematic, dynamics of bacterial abundance within a community comprising 3 bacteria (left, red) is approximated using *K* = 2 ECNs {*y*_*k*_ (*t*)} and corresponding loadings {**Φ**_*k*_} (middle). From the abundance data, EMBED identifies ECNs that are shared across subjects (right). The dynamics of abundances of individual bacteria are then approximated (green) using the inferred ECNs. **(B)** Average Kullback-Leibler divergence, averaged over the total duration, between observed microbial abundances and reduced dimensional reconstructions using EMBED and CTF using *K* = 5 components. Each line represents one subject. **(C)** Symmetric Kullback-Leibler divergence (Jensen-Shannon divergence) between two models learned from two different multinomial samplings of the same underlying ground truth microbiome trajectories generated using the multispecies Lotka-Volterra model across different sequencing depths. Error bars represent means and standard error of the means over 100 independent simulations. **(D)** Fraction of taxa that correlated with only one (left) and more than one ECN (right) obtained using EMBED, temporal components obtained using CTF, and temporal component obtained using singular value decomposition of the ***zθ*** matrix (Eq. 2). *p* values are calculated using the Wilcoxon signed rank test (*: *p* < 10^−9^).

In Eq. 2, *Z*_*tk*_ are time-specific latents that are shared by all OTUs and subjects, *θ*_*kos*_ are OTU- and subject-specific loadings that are shared across all time points, and Ω_*st*_ is the normalization constant. This low rank tensor factorization is a special case of the so-called Tucker decomposition^28^. The number of latents/loadings is chosen such that *K* ≪ *O,T* thereby achieving a reduced dimensional description of the data. These parameters are estimated using log-likelihood maximization.

The optimal values of the parameters depend on the initial conditions but are nonetheless related to each other via a linear transformation^27^. We therefore identify a unique and orthonormal representation for the latents by exploiting the dynamical nature of the data. The long-term stability of the microbiome is now well-established^16,17,29^. Therefore, we fit a model “return to normal” linear dynamical model to inferred latents:

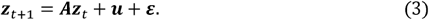

In Eq. 3, the matrix ***A*** is assumed to be symmetric, ***u*** are the baseline values, and the noise ***ε*** Gaussian distributed and uncorrelated. After diagonalizing the inferred interaction matrix (see **Supplementary Information**), ***A*** = ***v***^*T*^**Λ*v***, we find that the re-oriented latents, or the *ecological normal modes* (ECNs), ***y***_*t*_ = ***vz***_*t*_ fluctuate independently of each other

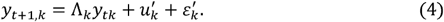

In Eq. (4) ***u***′ = ***vu***, and ***ε***′ = ***vε***. We redefine the corresponding loadings **Φ** = ***v***^*T*^***θ***. Notably, since ***vv***^*T*^ *= I*, this simultaneous transformation is a mere re-orientation of the latents and the loadings and does not change model predictions^27^. As we show in the **Supplementary Information**, the orthonormal ECNs are uniquely defined (SI Fig. 1) for a given dataset. We note that the actual dynamics of the latents are likely to be more complex than the linear model (Eq. 3). Yet, similar to normal mode analysis^26^, ECNs represent a re-orientation of the latents that uncovers the *unique* and *orthogonal* templates of microbial abundance fluctuations.

### EMBED accurately and robustly approximates microbiome abundance time series using dynamics on a lower dimensional manifold

Using EMBED, we approximated microbiome time series from several publicly available longitudinal data sets^10–14^ (see **Supplementary Information**). EMBED based reconstruction was highly accurate even when only a small number of ECNs was used (Fig. 1B, SI Fig. 2). In contrast, CTF (compositional tensor factorization), a recently developed dimensionality reduction method by Martino et al.^18,25^ had consistently larger reconstruction error (Fig. 1B, SI Table 1). EMBED-based reconstruction is also accurate for the time series of individual bacterial taxa. The taxon-specific Pearson correlation coefficient between the reconstruction and the data averaged across taxa and datasets for humans was *r* = 0.72 ± 0.06 and 0.84 ± 0.06 for mice, compared to an average correlation of *r* = 0.52 ± 0.04 for humans and *r* = 0.68 ± 0.05 for mice for the CTF-based reconstruction. Collectively, these results show that microbiome dynamics are surprisingly low dimensional and EMBED can accurately approximate these dynamics by embedding them in a lower dimensional manifold.

The probabilistic formulation of EMBED naturally accounts for spurious abundance variability arising from sampling noise. To test the robustness of EMBED to sampling noise, we generated ground truth trajectories using the multispecies Lotka-Volterra model^30^ with competitive interactions^31,32^ (see **Supplementary Information**). Using different sequencing depths, two sets of read counts were sampled using the same ground truth abundances. EMBED (and CTF) was used to model the observed read counts. The more robust the inference is to sampling noise, the better will be the agreement between the two inferred models. Indeed, as seen in Fig. 1C (SI Fig. 3), EMBED-based reconstruction of abundance time series was internally consistent and robust to sequencing noise.

The re-orientation ***z* → *y*** of latents using a dynamical model (Eq. 3 and 4) allows us to identify independent directions of significant collective dynamics in the microbiome without changing the accuracy of model predictions. In contrast, any other orthogonal decomposition of the microbiome time series that does not explicitly model dynamics is likely to result in a latent space description that involves a mixture of independent modes. To test the dynamical independence of ECNs (Fig. 1D), we generated 100 microbiome time series using the competitive multispecies Lotka-Volterra model as above. Each time series was approximated using EMBED using *K* = 5 ECNs (SI Fig, 4). As with Fig. 1B, EMBED-based approximation of the microbiome time series was significantly accurate (mean average KL divergence 0.0143 vs 0.0182 for CTF, Wilcoxon signed rank test *p* = 3.9×10^−18^). We correlated the inferred ECNs with time series of abundances of individual taxa. Correlations that were above a 5% FDR using the Benjamini-Hochberg procedure were deemed significant. Importantly, on an average, 57% of OTUs correlated with only one ECN while only 24% of OTUs correlated with two or more ECNs. This suggests that ECNs represent modes of independent collective dynamics rather than a mixture of several modes. In contrast, 44% of OTUs correlated with only one component obtained using CTF (Wilcoxon signed rank test *p* = 5.1×10^−8^) and 44% OTUs correlated with two or more components (Wilcoxon signed rank test *p* = 3.7×10^−12^). Notably, the specificity of taxon-ECN correlations was not due to the accuracy of the EMBED-based reconstruction. To test this, we performed SVD on the ***zθ*** matrix prior to the re-orientation step (Eqs. 3 and 4 above) to obtain orthonormal latents ***y***_*SVD*_ that *did not* take into account the longitudinal nature of the data. We found that statistics of correlations of individual bacterial taxa with ***y***_*SVD*_ were indistinguishable from CTF and significantly different compared to ECNs (Fig. 1D). These analyses underscore the importance of dynamics-based re-orientation of the latents in EMBED in identifying independent modes of significant collective abundance changes.

### Effect of dietary oscillations on the gut microbiome

Host diet has been shown to be a major factor influencing gut bacterial dynamics^13,33^ but in a subject specific manner^34^. We applied EMBED to the data collected by Carmody et. al.^13^ to better understand bacterial abundance changes in response to highly controlled dietary perturbations. Briefly, the diets of five individually housed mice were alternated every ∼3 days between a low-fat, plant-polysaccharide diet (LFPP) and a high-fat, high-sugar diet (HFHS). Daily fecal samples were collected for over a month (SI Fig. 5).

Using *K* = 5 ECNs, EMBED obtained a lower dimensional time series approximation that reconstructed the original data with great accuracy (average taxa Pearson correlation coefficient *r* = 0.75±0.18, average community Pearson correlation coefficient, *r* = 0.98± 0.003) (SI Fig. 6). The first ECN *y*_1_(*t*) represented a relatively constant abundance throughout the entire time series (Fig. 2A). Moreover, the corresponding loading vector **Φ**_1_ showed a significant correlation to the average individual OTU abundance across time. (Average Spearman correlation coefficient across subjects, *r* = −0.86±0.06, SI Fig. 7), suggesting that despite large-scale, cyclic dietary changes, gut bacterial abundances in the community tended to fluctuate around a constant average abundance.

**Figure 2.**
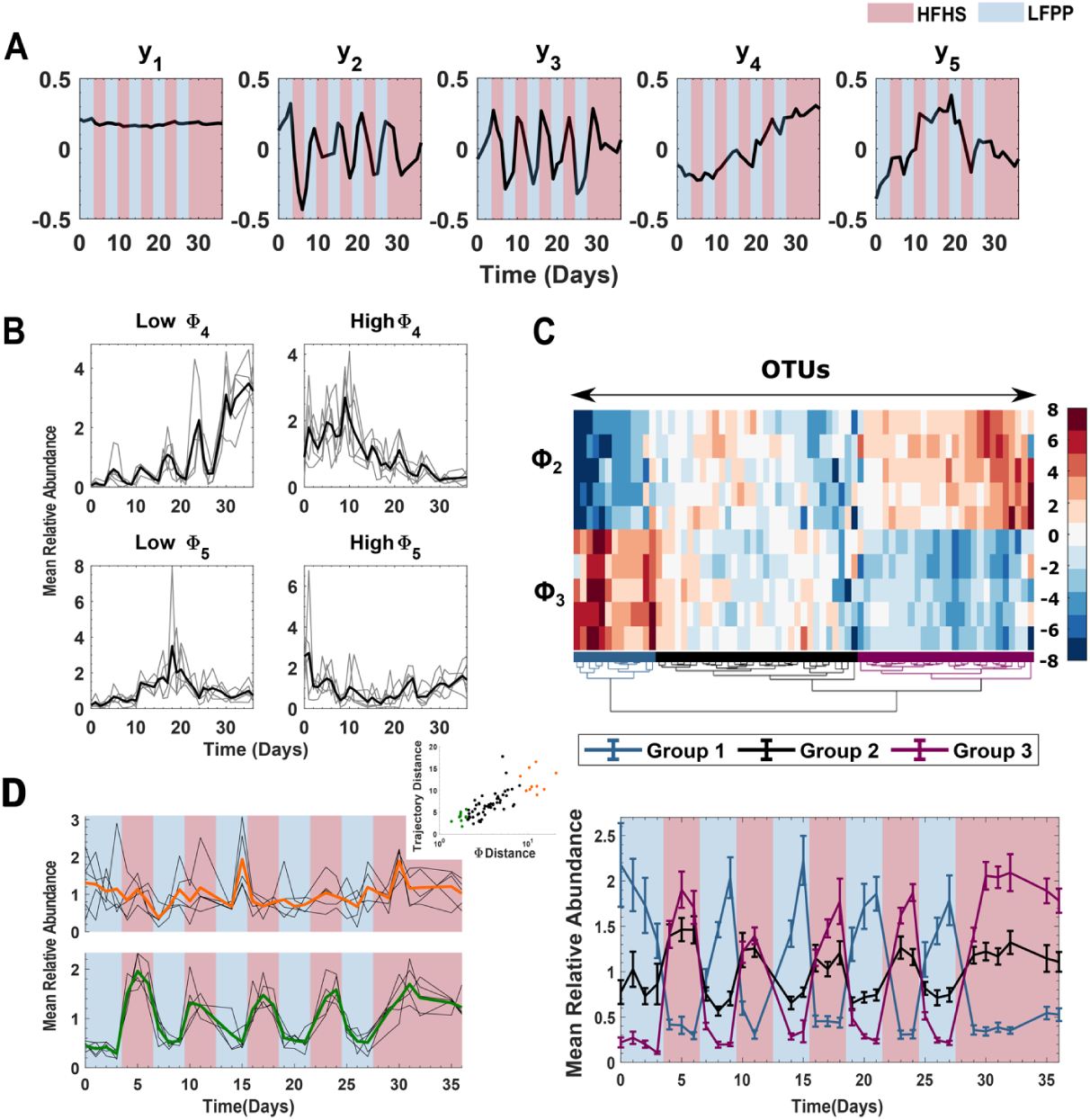
The effect of dietary oscillations on microbiome dynamics. **(A)** Temporal profiles of the five inferred ECNs. Blue and red panels show periods of time of administered LFPP and HFHS diets respectively. (**B**) (Top) The average abundances of five OTUs with the most negative and the most positive **Φ**_4_ values. (Bottom) The average abundances of five OTUs with the most negative and the most positive **Φ**_5_ values. For each subject, the abundances of the identified OTUs were first mean-normalized for each OTU, then averaged across the OTUs (faint lines). The bold lines show abundances averaged across all subjects. (**C)** (Top) A hierarchical clustering of OTUs using the two oscillatory loadings **Φ**_2_ and **Φ**_3_ identifies three major groups of OTUs (colored). (Bottom) Mean relative abundance of OTUs in the three groups using the same colors as the top panel. The abundances were first mean-normalized on a per OTU basis, then averaged across subjects for each OTU, and then averaged across all OTUs in any given group. The error bars represent standard errors of mean estimated using the considered OTUs. (**D**) Abundance variation in top 10 OTUs that exhibit universal dynamics (green) and top 10 OTUs that show subject-specific dynamics (orange) as identified by the average subject-to-subject variability in OTU-specific **Φ** loadings (inset).

In contrast, ECNs *y*_2_(*t*) and *y*_3_(t) collectively captured the cyclic nature of dietary oscillations, confirming that the murine diet rapidly and reproducibly alters abundance dynamics even at the individual OTU level. To identify OTUs whose oscillatory dynamics were similar across subjects, we clustered the loadings **Φ**_2_ and **Φ**_3_ of individual OTUs on ECNs *y*_2_(*t*) and *y*_3_(*t*). We found that bacteria in the community largely clustered into three groups (Fig. 2C); those whose abundances increased with the LFPP diet (blue, group 1), and those whose abundances increased with the HFHS diet to different extents (black and magenta, groups 2 & 3). In keeping with recent studies^35–37^, we found that the genera *Saccharicrinis*, members of the Bacteroidetes phylum, were significantly enriched in group 1 (5 out of 13 compared to 7 out of 73, hypergeometric test, *p* = 0.0015) consistent with the notion that bacteria belonging to this genera are able to degrade plant polysaccharides and utilize the metabolic byproducts present in the LFPP diet.

Unexpectedly, we found two ECNs *y*_4_(*t*) and *y*_5_(*t*) that represented profound non-oscillatory behavior in abundance fluctuations. *y*_4_(*t*) represented an overall drift in abundance over the time series and *y*_5_(*t*) represented a U-shaped recovery. The loadings corresponding to these two modes the were significantly correlated across subjects (Spearman correlation coefficient *r* = 0.37±0.16, averaged across mice). The top 5 OTUs with most negative and positive loadings **Φ**_4_ (omitting OTUs that were also in the top 5 negative/positive for loadings **Φ**_5_ experienced a significant, irreversible increase and decrease throughout the time course of the experiment respectively (Fig. 2B top). Thus, while the dynamics of most gut bacteria in this community exhibit rapid and reversible changes in response to dietary oscillations, there exist certain bacteria that exhibit irreversible changes over time. This concept of *hysteresis* has been explored previously in the gut microbiome^13^, but the underlying mechanisms likely warrant continued investigation. In contrast, the top 5 OTUs with most negative and positive loadings **Φ**_5_ (omitting OTUs that were also in the top 5 negative/positive for loadings **Φ**_4_) experienced an inverted U-shaped and a U-shaped abundance profile (Fig. 2B bottom). Interestingly, OTUs that exhibited these non-oscillatory behaviors differed significantly from subject-to-subject (SI Table 2, SI Fig. 5). This *ecological equivalence*^38^ is reminiscent of the macroecological laws recently reported by Ji et al.^16^ that were shared across different host organisms but were exhibited by different bacterial taxa.

EMBED can identify OTUs that exhibit universal dynamics and those that exhibit subject-specific behavior. Each OTU within each subject-specific ecosystem is characterized by a *K*-dimensional vector of loadings corresponding to the *K* ECNs. OTUs whose loading vectors are similar across all subjects have similar dynamics across subjects and vice-versa for OTUs with different loading vectors. To identify these universal and subject specific OTUs, we computed the average distance across all pairs of subjects of the OTU specific loadings vectors. This average distance also correlated strongly with the average distance of the subject specific OTU abundance trajectories (inset of Fig. 2D). In Fig. 2D, we plot the average abundance of 10 OTUs with the most similar **Φ** loadings (bottom) and the 10 most dissimilar **Φ** loadings (top). The black lines show the OTU-averaged abundances for individual subjects and the colored bold lines (green and orange) show the average across subjects. As seen in Fig. 2D, the top 10 OTUs whose dynamics were similar across all subjects strongly preferred the HFHS diet. Notably, these OTUs are overrepresented by the genus *Oscillibacter* (4 out of 10 compared to 5 out of 73, Hypergeometric test *p* = 9×10^−4^). Interestingly, this overrepresentation was observed only at the genus and the family level and was *not* observed at higher taxonomic classifications (SI Table 3). Moreover, no other genus or family were overrepresented. This strongly suggests a specific genus level preference to high fat high sugar diet in the genus *Oscillibacter* that can override subject-specific ecosystem parameters. Notably, *Oscillibacter* are known to prefer high fat^39^ as well as high sugar diets^40^. Future work is needed to further establish the mechanistic connection between *Oscillibacter* and HFHS diets.

### ECNs identify modes of recovery of bacteria under antibiotic action

Broad-spectrum oral antibiotics have significant effects on the gut flora both during and after administration. Specifically, microbiome abundance dynamics following antibiotic administration can potentially exhibit a combination of several typical behaviors which may reflect different survival strategies^7,9,11,41^. These include quick recovery following removal of antibiotic, slow but partial recovery, and one-time changes followed by resilience to repeat antibiotic treatment. The temporal variation in abundances of any bacteria could be a combination of these typical behaviors. Moreover, given that the gut ecosystems differ across different hosts, the response of specific bacteria to the same antibiotic treatment could vary from host to host. To better parse the major modes of gut bacterial dynamics associated with antibiotic administration, we analyzed the data collected by Ng et al.^10^. Briefly, six mice were given the antibiotic ciprofloxacin in two regimens (days 1-4 and days 14-18) and fecal microbiome samples were collected daily over a period of ∼ 30 days (SI Fig. 8).

We found that a very small number ECNs was sufficient to capture the data with significant accuracy (average taxa Pearson correlation coefficient, average community Pearson correlation coefficient,) (SI Fig. 6). As shown in Fig. 3A and consistent with the diet analysis, we found that ECN was relatively stable throughout the study and the corresponding loading vector was strongly correlated with the mean OTU abundance over time (Spearman correlation coefficient) (SI Fig. 7). We found the remaining several ECNs to follow broad classes of behaviors in response to periods of stress. Indeed, ECNs, appeared to represent an inelastic one-time change followed by a relatively stable response. ECN, represented the opposite, it responded to the antibiotic treatment the second time but not the first time. In contrast, ECN represented *elastic* changes in the microbiome, potentially representing abundances reproducibly decreasing (or increasing) with the action of the antibiotic but quickly bouncing back to pre-antibiotic levels when it was withdrawn.

**Figure 3.**
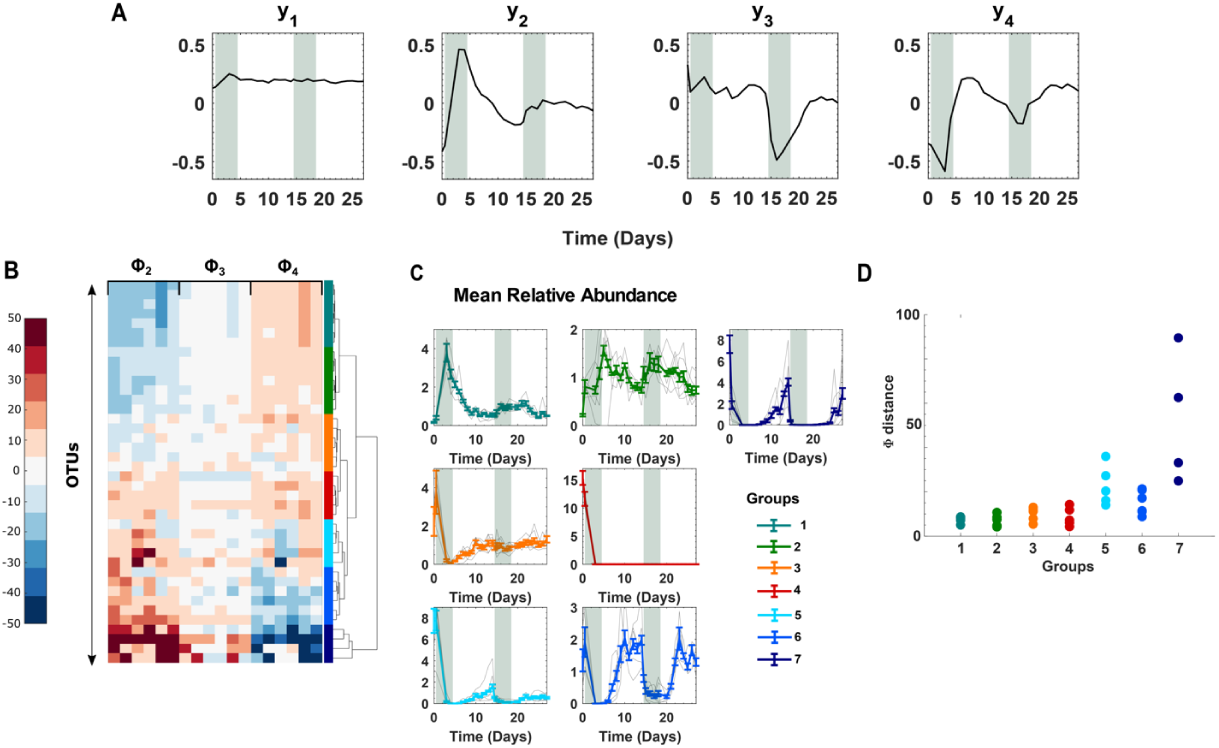
Effect of antibiotic treatment on the gut microbiome. **(A)** ECNs describe the microbiome of mice on antibiotics. The shaded region indicates the first and second doses of ciprofloxacin. **(B)** A hierarchical clustering of OTUs using loadings except for. 7 major groups of OTUs with similar dynamical responses are identified from the clustering. **(C)** In every group and for each subject, the abundances of the identified OTUs were first mean-normalized at the OTU level. The faint lines represent subject-specific average over OTUs. The bold lines represent average across subjects. Error bars represent standard errors of mean estimated using the considered OTUs. **(D)** Average subject-to-subject variability in OTU-specific **Φ** loadings for the 7 identified groups.

These salient dynamical features were captured when we clustered the OTUs using the loadings **Φ**_2_ − **Φ**_4_ (panel B), which identified seven major groups of OTUs with distinct dynamical behaviors (Figure 3B and C). Interestingly, while some of the groups simply reflected behaviors of individual ECNs, others could be understood according to their relative contributions across multiple ECNs. For example, the behavior of OTUs in groups 1 and 3 aligned with ECN *y*_2_(*t*), albeit with opposing trends. Group 1 OTUs flourished during the first antibiotic treatment but the second treatment did not elicit a similar response. In contrast, OTUs in group 3 diminished in their abundance after the first antibiotic treatment but were resistant to subsequent antibiotic action.

OTUs in groups 2, 5, 6, and 7 displayed highly elastic dynamics in response to both periods of antibiotic administration. Group 2 OTUs overrepresented by the genus *Akkermansia* (all 2 out of 41 OTUs are in Group 2, Hypergeometric test *p* = 0.026) flourished during the antibiotic treatment but decreased their abundance in a reversible manner when antibiotics were withdrawn. Notably, species from this genus are known to be rare in the human gut but only colonize it following treatment with broad spectrum antibiotics including ciprofloxacin^42^. OTUs in groups 5, 6, and 7 in contrast diminished their abundance in the presence of antibiotics in a reversible manner. Group 6 was overrepresented by the genus *Blautia* (3 out of 6 compared to 5 out of 41, Hypergeometric test *p* = 0.017), while group 7 was overrepresented by the genus *Aestuariispira* (all 2 out of 41 OTUs are in Group 7, Hypergeometric test *p* = 0.0073). Finally, group 4 comprised OTUs that were exquisitely sensitive to initial antibiotic administration, whose abundance did not make any meaningful recovery. These OTUs were overrepresented in the genus *Coprobacter* (2 out of 5 compared to 3 out of 41, Hypergeometric test *p* = 0.035). These specific associations need to be further investigated.

Notably, OTUs in groups 5 and 7, groups that represent slower and partial recovery compared to OTUs group 6, exhibited significant subject-to-subject variability as quantified by both the average subject-to-subject variability in OTU-specific **Φ** loadings (Fig. 3D) and the subject-to-subject variability in OTU-specific abundance trajectories (SI Fig. 8). While these OTUs exhibited qualitative dynamics of recovery across all subjects (SI Fig. 8), the time course and the extent of recovery varied from subject-to-subject. These findings are corroborated by recent studies that show imperfect and subject-specific recovery of bacterial abundances following antibiotic treatment^11,43–45^.

## Discussion

Bacteria in host-associated microbiomes live in complex ecological communities governed by competitive and cooperative interactions, and a constantly changing environment. Extensive spatial and temporal variability and coordinate changes in abundances in response to environmental perturbations are a hallmark of these communities. Dimensionality reduction can leverage these fluctuations, but its use towards understanding microbiome dynamics has thus far been limited.

In this work, we presented EMBED, a novel dimensionality reduction approach specifically tailored to identify the *ecological normal modes* in the dynamics of bacterial communities that are shared across subjects undergoing identical environmental perturbations. ECNs can be viewed as dynamical templates along which the trajectories of individual bacteria within individual host ecosystems can be decomposed. Identified ECNs shed insight into the underlying structure of bacterial community dynamics. By applying EMBED to several times series data sets representing major ecological perturbations, we identified immediate and reversible changes to the gut community in response to these stimuli. However, EMBED also identified more subtle, longer-term, and perhaps irreversible changes to specific members of the community, the mechanisms and consequences of which would be interesting to pursue further.

One key parameter in EMBED is the number of components *K*. A large *K* will necessarily fit the data better, potentially fitting to noise and unimportant idiosyncrasies in the data. How do we decide the appropriate number of components? EMBED is a probabilistic model and information theoretic criteria^46^ could be used to identify the correct number of components. These criteria seek a balance between increase in number of parameters and the accuracy of fit to data (likelihood). We note that the total likelihood of the data in our model is linearly proportional to the sequencing depth. However, the reported sequencing depth is typically over-inflated compared to the true nucleotide capture probability of the experiments leading to an inflated estimate of the total likelihood. This issue has been well discussed in single cell RNA sequencing (see for example^47^). One approach to solve this in the context of the microbiome to obtain technical repeats which can in turn allow us to estimate the true technical noise.

While EMBED was specifically developed to study microbiomes, it reflects a more generalizable framework that can easily be applied to other types of longitudinal sequencing data as well. We therefore expect that EMBED will be a significant tool in the analysis of dynamics of high dimensional sequencing data beyond the microbiome.

