## Supplementary text and figure captions for "EMBED: Essential Microbiome Dynamics, a dimensionality reduction approach for longitudinal microbiome studies"

**Supplementary Information**

1. **Inference of ECNs from longitudinal data**

We consider that abundance of $O$ bacterial operational taxonomic units (OTUs) are measured over a period of $T$ days in $S$ subjects. We model the read counts $n_{os}(t)$ of OTUs $"o"$ on any given day $t$ in subject $s$ as a multinomial distribution. The likelihood of observing the data is given by

$$L=\prod_{s,t} \frac{N_{s}\left( t \right)!}{\prod_{o} n_{os}\left( t \right)!}\prod_{o} q_{os}\left( t \right)^{n_{os}\left( t \right)} (S1)$$

where $N_{s}\left( t \right)=\sum_{o} n_{os}(t)$ is the total read count on a given day and $q_{os}\left( t \right)$ are the underlying propensities for individual OTUs. We model these propensities using the exponential Gibbs-Boltzmann distribution which allows us to capture large variations in OTU abundances^1^

$$q_{os}\left( t \right)=\frac{1}{\Omega_{st}}\exp\left( -\sum_{k=1}^{K} z_{tk}\theta_{kos} \right) (S2)$$

where $z_{tk}$ are time-specific latents that are shared by all OTUs and subjects, and $\theta_{kos}$ are OTU-and subject-specific loadings that are shared across all time points. The number $K$ of latents/loadings is chosen such that $K\ll O, T$ thereby achieving a lower dimensional description of the time series data. We obtain the$z$s and the $\theta$s using the maximum likelihood approach.

To that end, we write down the log-likelihood of the data:

$$ln=const.+\sum_{t,o,s} n_{os}\left( t \right)\log q_{os}(t). (S3)$$

The constant term of the likelihood does not depend on the parameters and can thus be omitted in likelihood maximization. Simplifying using Eq. S1 and S2, we have

$$ln=-\sum_{t,o,s,k} {N_{s}\left( t \right)x}_{os}\left( t \right)z_{tk}\theta_{kos}-\sum_{t,s} \log\Omega_{st} (S4)$$

Here $x_{os}\left( t \right)=n_{os}(t)/N_{s}(t)$ is the relative abundance of OTU $o$ at time $t$. We obtain the gradients

$$\frac{\partial ln}{\partial z_{tk}}=-\sum_{o,s} N_{s}(t)\left( x_{os}\left( t \right)-q_{os}\left( t \right) \right)\theta_{kos} \mathrm{and} (S5)$$

$$\frac{\partial ln}{\partial\theta_{kos}}=-\sum_{t} N_{s}\left( t \right)z_{tk}\left( x_{os}\left( t \right)-q_{os}\left( t \right) \right) (S6)$$

We use gradient ascent algorithm to find the latents and the loadings that maximize the likelihood.

For a given $K$, using the microbiome data $x_{os}\left( t \right)$ and starting from random initialization, we first simultaneously infer the latents $z_{tk}$ and the features $\Theta_{kos}.$ We observe that the $T\times K$ matrix $\boldsymbol{z}$ of latents can be multiplied by an invertible matrix $\boldsymbol{B}$ $\left( \boldsymbol{z\to zB} \right)$ and the corresponding matrix $K\times O\times S$ matrix of features can be multiplied by the inverse $\boldsymbol{B}^{\boldsymbol{-1}}$ $\left( \boldsymbol{\Theta}\boldsymbol{\to}\boldsymbol{B}^{\boldsymbol{-1}}\boldsymbol{\Theta} \right)$ and the abundance predictions from the model do not change. Therefore, we use the Gram-Schmidt procedure to orthogonalize the matrix of latents such that $\boldsymbol{z\to}\boldsymbol{z}^{\boldsymbol{'}}$where ${\boldsymbol{z}^{\boldsymbol{'}}}^{T}\boldsymbol{z}^{\boldsymbol{'}}\boldsymbol{=}\boldsymbol{I}_{\boldsymbol{K}}$ is an identity matrix. For any matrix of latents $\boldsymbol{z}$, the matrix multiplier $\boldsymbol{B}$ that leads to the orthonormal transformation can be found by solving the equation $\boldsymbol{B}^{\boldsymbol{T}}\left( \boldsymbol{z}^{\boldsymbol{T}}\boldsymbol{z} \right)\boldsymbol{B=}\boldsymbol{I}_{\boldsymbol{K}}\boldsymbol{.}$ Once $\boldsymbol{B}$ is identified, we also transform the $\boldsymbol{\Theta}$ matrix ($\boldsymbol{\Theta\to}\boldsymbol{\Theta}^{\boldsymbol{'}}\boldsymbol{=}\boldsymbol{B}^{\boldsymbol{-1}}\boldsymbol{\Theta}\boldsymbol{)}$. At the end of this procedure, we end up with orthonormal latents $\boldsymbol{z}^{\boldsymbol{'}}$and corresponding features $\boldsymbol{\Theta}\boldsymbol{'}$ that correspond to the same abundances as $\boldsymbol{z}$ and $\boldsymbol{\Theta}$. For the sake of simplicity of notation, we drop the primes.

Next, we model the dynamics of the orthonormal latents using a linear dynamical system:

$$z_{t+1,k}=\sum_{k^{'}} A_{kk^{'}}z_{{tk}^{'}}+u_{k}+\eta_{k}(t) (S7)$$

where we assume that $A_{kk^{'}}=A_{k^{'}k}$ and $\eta_{k}(t)$ are Gaussian distributed uncorrelated noise vectors: ${\langle\eta}_{k}\left( t_{1} \right)n_{k^{'}}\left( t_{2} \right)\rangle=\delta_{12}\delta_{kk^{'}}$ where $\delta_{ab}$ is the Kronecker delta function. Our task is to find the interaction matrix $\boldsymbol{A}$ and the vector $\boldsymbol{u}$ that fits this model. We achieve this using squared error minimization. We write

$$E\left( \boldsymbol{A,u} \right)=\sum_{t} \left( z_{tk}-z_{tk}^{pred} \right)^{2} (S8)$$

where $z_{tk}$ is the inferred latent and $z_{tk}^{pred}$ is the corresponding prediction using $z_{t-1,k}$ and Eq. S7. We restrict the summation only over time points $t$ such that measurements are available for time points $t$ and $t-1$. The squared error is minimized using a simulated annealing approach. Once the matrix $\boldsymbol{A}$ is identified, we transform the orthonormal latents $z_{tk}$ into ecological normal modes $y_{tk}$ as described in the manuscript.

The scripts for obtaining ECNs $\boldsymbol{y}$ and corresponding loadings $\boldsymbol{\Phi}$ from read count data can be found at: https://github.com/mayar-shahin/EMBED.

2. **Generating *in silico* data using the multispecies Lotka-Volterra model with competitive interactions.**

The multispecies Lotka-Volterra model is one of the most popular models to study microbiome dynamics^2^. We use a discrete time multispecies Lotka-Volterra model introduced by Kilpatric and Ives^3^. Briefly, the abundances of species are governed by the following equation:

$$x_{i}\left( t+1 \right)=x_{i}\left( t \right)\exp\left[ r_{i}\left( 1-\frac{x_{i}\left( t \right)+\sum_{j} \alpha_{ij}x_{j}(t)}{K_{i}} \right)+\epsilon_{i}(t) \right]. (S9)$$

In Eq. S9, $x_{i}\left( t \right)$ is the abundance of species $"i"$ at time $t$, $r_{i}$ is the species-specific growth rate, $K_{i}$ is the species-specific carrying capacity, $\alpha_{ij}$ is the pairwise interaction matrix, and $\epsilon_{i}$ is uncorrelated Gaussian distributed noise.

For our simulations, we chose a system with $O_{total}=200$ organisms. The matrix of interactions $\boldsymbol{\alpha}$ was assumed to be symmetric and had only positive entries reflecting the highly competitive nature of interactions amongst bacteria in microbiome communities^2,4^. Moreover, we assumed that the matrix of interactions was sparse (only 10% of the entries were non-zero). The non-zero entries of the interaction matrix were chosen uniformly between 0 and 0.1. The species intrinsic growth rates were chosen from a uniform random distribution between 0 and 0.05. The carrying capacities were chosen from a uniform random distribution between 0 and 1. Finally, the standard deviation of the noise was set to 0.2. A community of $O_{total}$ organisms was initialized from random initial abundances between 0 and 1 and propagated for 300 time steps. Only the last 30 time steps were used for further analysis. The mean abundances of all species in the last 30 time steps were calculated. The species whose mean abundances were less than 0.1% were clubbed together as a single species. This protocol typically led to an ecosystem with 50-100 species. Finally, the species abundances were normalized and then sampled using a multinomial distribution with sequencing depth of 5000. This data was used for further analysis.

3. **Obtaining the microbiome time series from sequencing data**

**Murine gut microbiome response to oscillating diet.** We downloaded the microbiome abundance time series data on mice fed an alternating diet of high fat high sugar chow (HFHS) and low-fat plant polysaccharide chow (LFPP) from Carmody et al.^5^ as described previously^6^. Each mouse that was subjected to an oscillatory diet was treated separately. Based on our previous work on technical noise in 16s measurements, we only analyzed OTUs with mean abundances > 0.1%^7^ averaged across all time points and mice. On every day, the abundances of the rest of the OTUs were lumped together in a single meta-species.

**Murine gut microbiome response to antibiotics.** We downloaded microbiome abundance data from Ng et al.^8^. We focused on the data where mice were administered the antibiotic ciprofloxacin. Out of the 10 cages in which the mice were housed, we omitted data from cages 2, 4, 5, and 8 where many time points were missing. As above, we analyzed OTUs with mean abundance > 0.1% and combined the rest of the OTUs in a meta-species.

4. **Performing CTF**

We downloaded publicly available data sets from several studies^5,8–11^. Each data set comprised microbiome abundance tables for multiple subjects, see SI Table 1. We use the package released by Martino et al.^12,13^ (<https://github.com/biocore/gemelli>) to perform compositional tensor factorization. To test how well the dimensionality reduced version capture the data, we calculate an approximate reconstruction of the abundance time series using the first *K* singular values. The resulting approximation was re-exponentiated and then normalized. We then calculated the KL-divergence with the true abundances.

**Supplementary Figures**

**Supplementary Figure 1**. Plot showing that the ecological normal modes (ECNs) inferred using EMBED are unique (up to a sign) for the two data sets considered in detail in this manuscript. The x- and the y-axis represent the ECNs inferred in two independent runs.

**Supplementary Figure 2**. (**A**) KL divergence, averaged over the duration of the study, for reconstruction of abundance time series using EMBED and CTF and $K=3,4,5$ components. (**B**) Scatter plot of approximated abundances using EMBED vs. measured abundances for $K=3,4,5.$

**Supplementary Figure 3.** Fig. 1C for $K=3,4,5.$

**Supplementary Figure 4.** Fig. 1D for $K=3,4,5.$

**Supplementary Figure 5.** Abundance time series of individual OTUs in the diet oscillation study. The gray lines represent abundances in individual subjects. The dark lines represent averages over subjects. The colors represent the cluster identities in main text Figure 2. HFHS: High Fat High Sugar diet and LFPP: Low Fat Plat Polysaccharide diet.

**Supplementary Figure 6.** Accuracy of reconstruction of abundance time series for the diet and the antibiotic study.

**Supplementary Figure 7.** Correlation of average OTU abundance (averaged over time) with the loading $\boldsymbol{\Phi}_{1}$ that corresponds with the constant ECN.

**Supplementary Figure 8.** Abundance time series of individual OTUs in the antibiotics-treatment study. The gray lines represent abundances in individual subjects. The dark lines represent averages over subjects. The colors represent the cluster identities in main text Figure 3. The gray bars represent the duration of time when the antibiotic was administered.
