## Supplementary figures and images for "EMBED: Essential Microbiome Dynamics, a dimensionality reduction approach for longitudinal microbiome studies"

### SI Figure 1

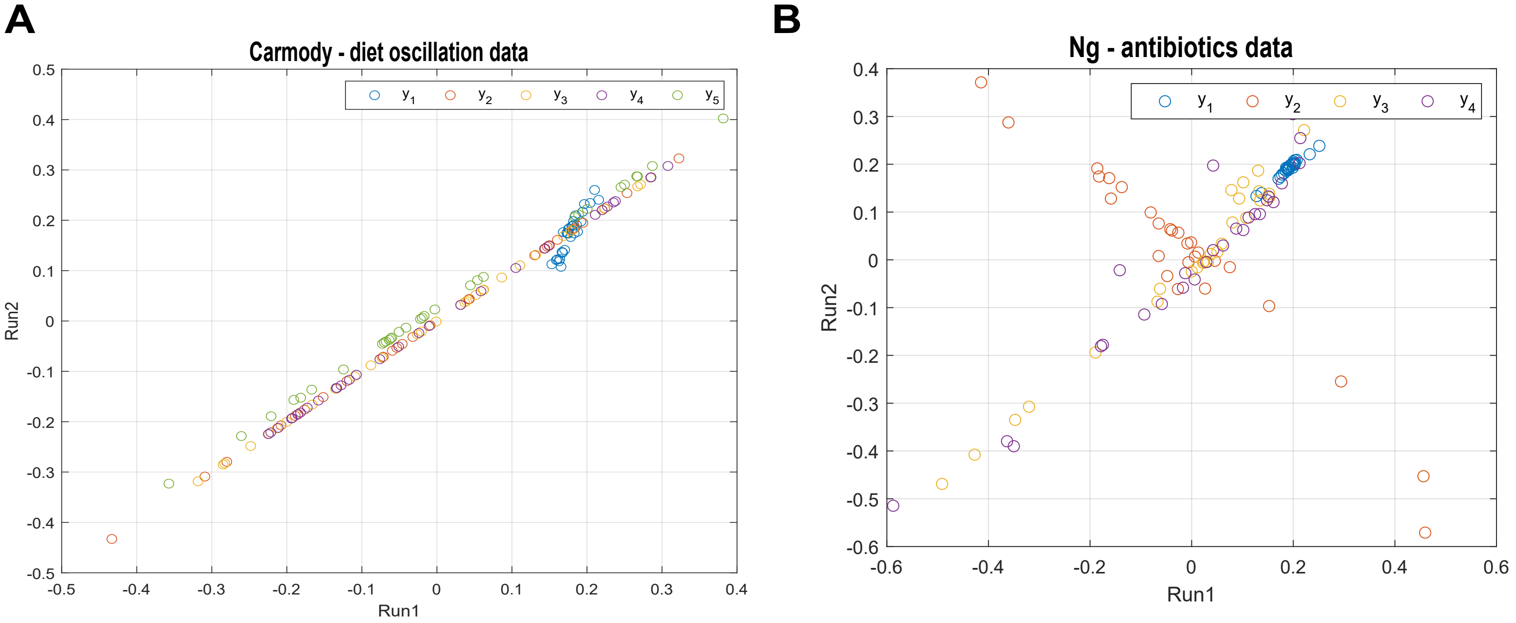

### SI Figure 2

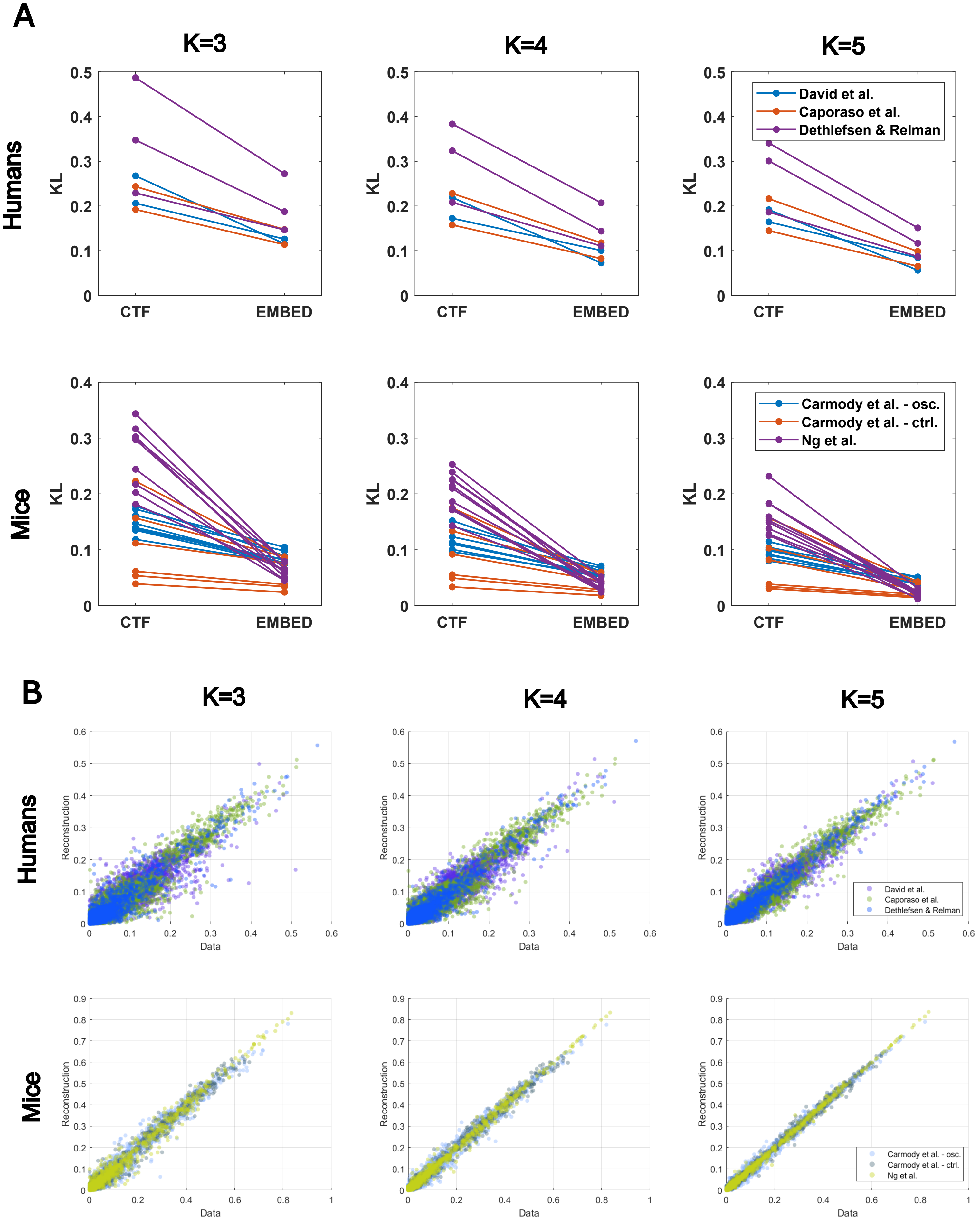

### SI Figure 3

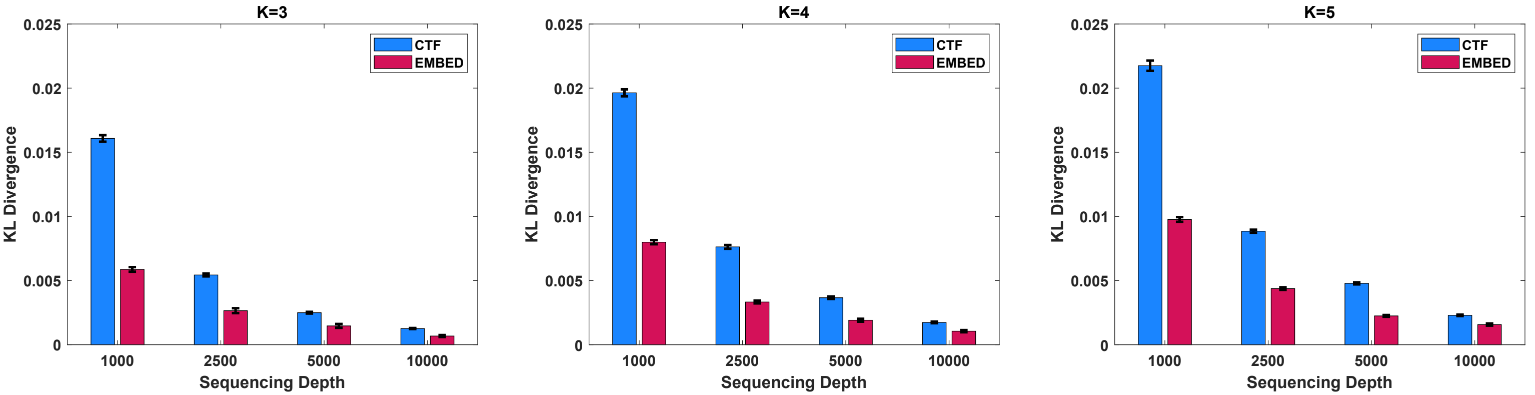

### SI Figure 4

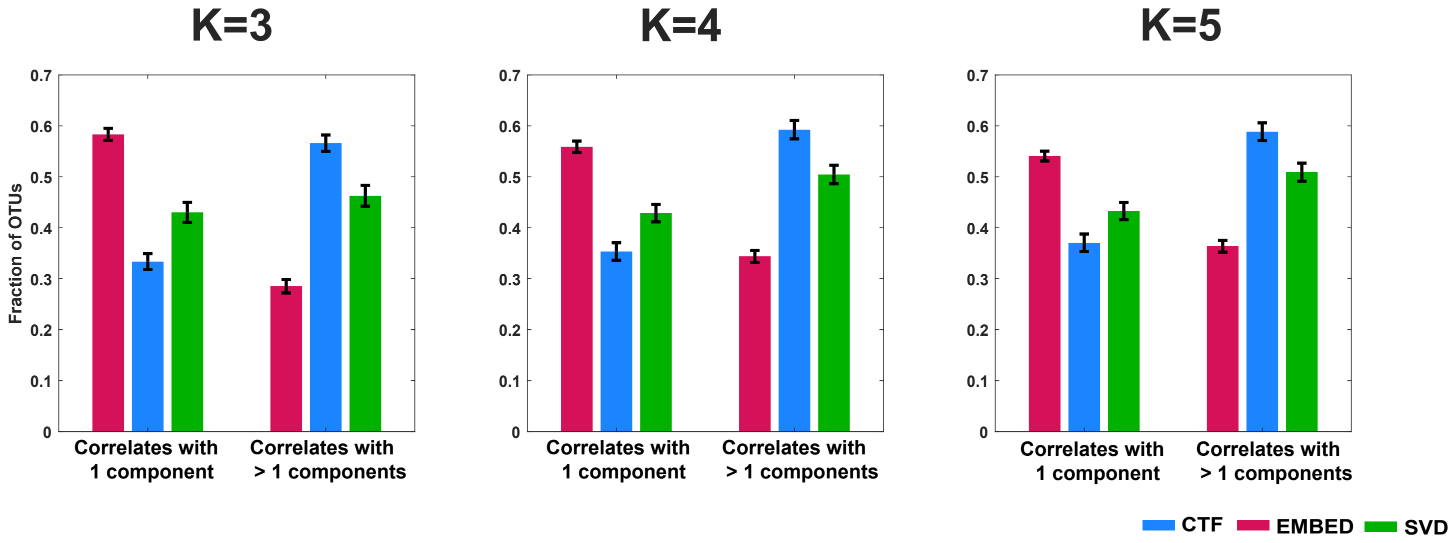

### SI Figure 5

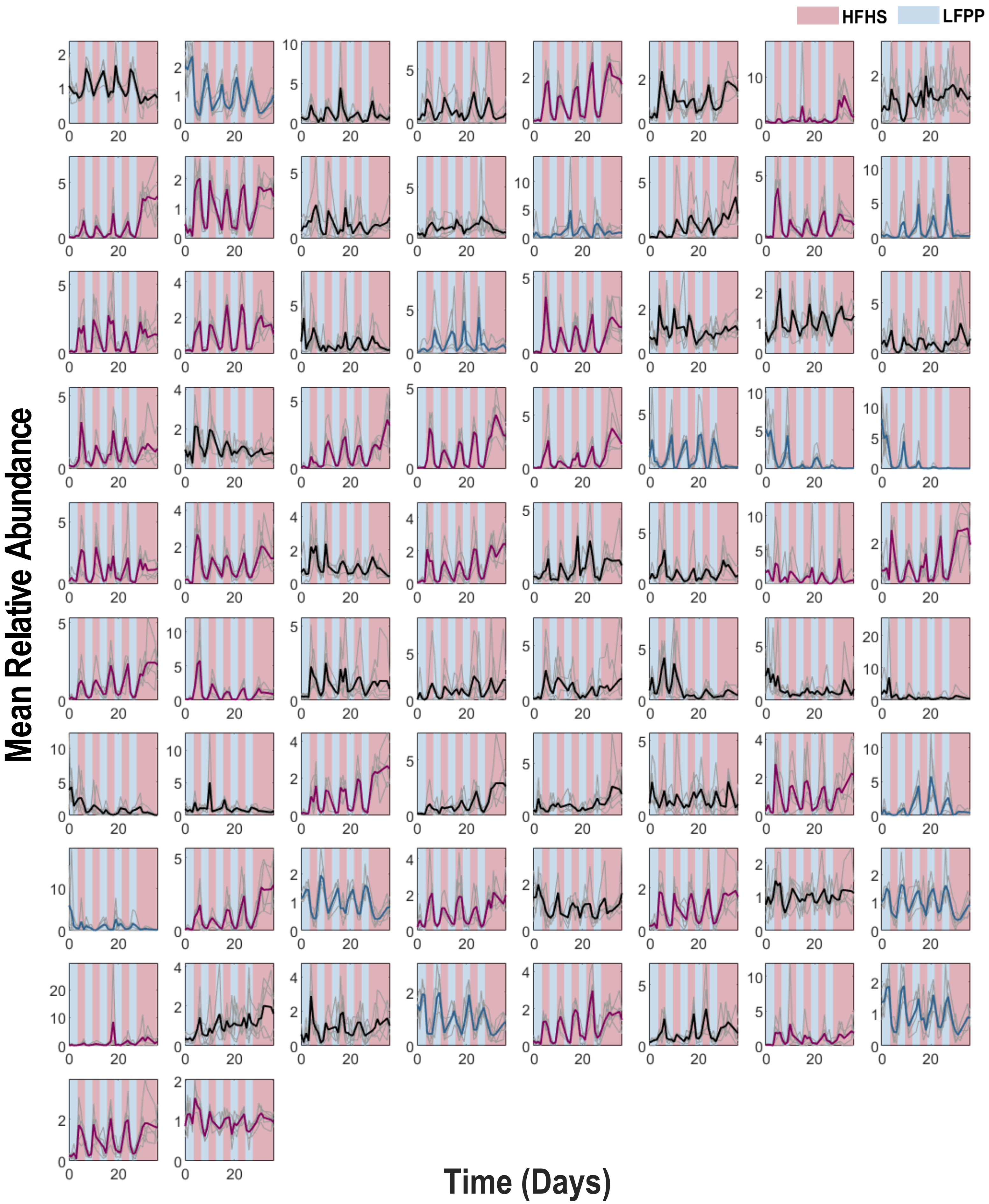

### SI Figure 6

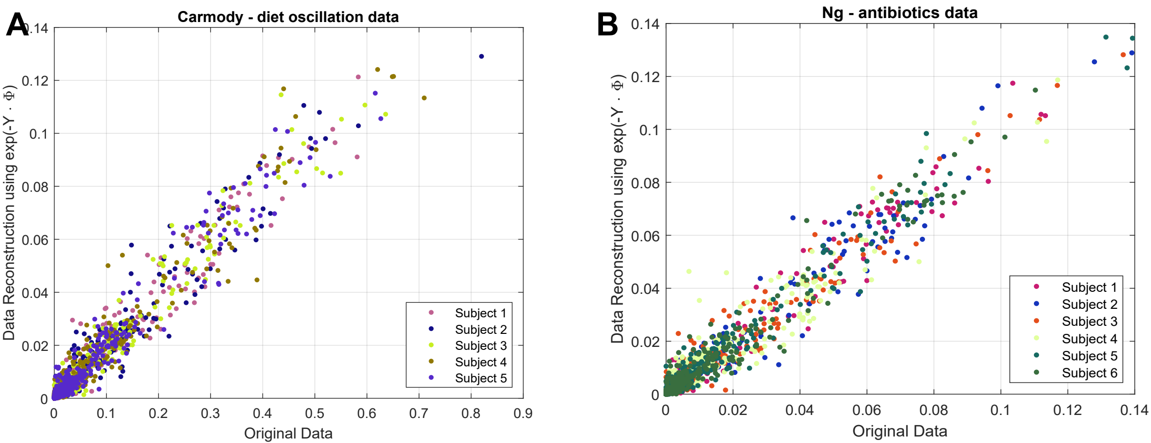

### SI Figure 7

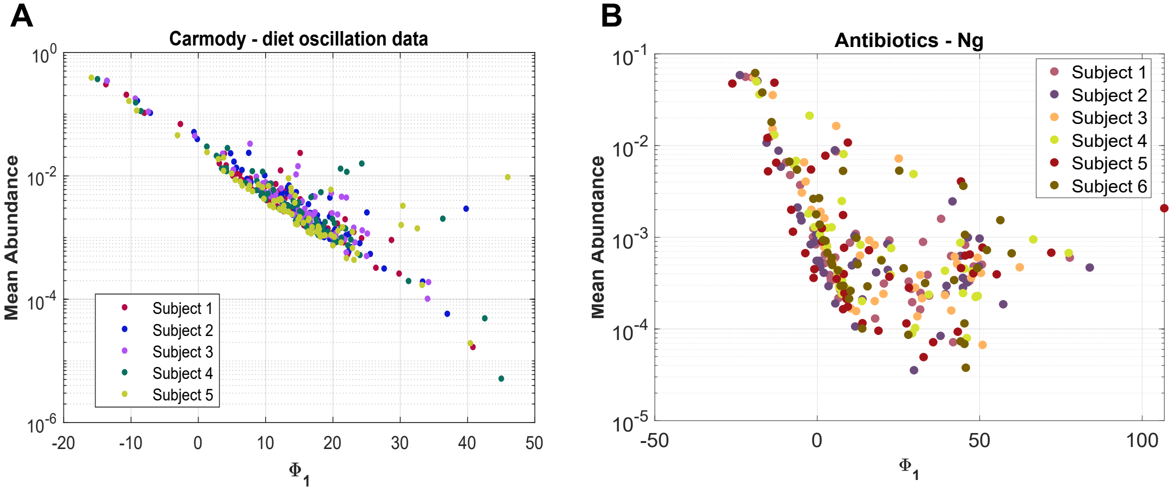

### SI Figure 8

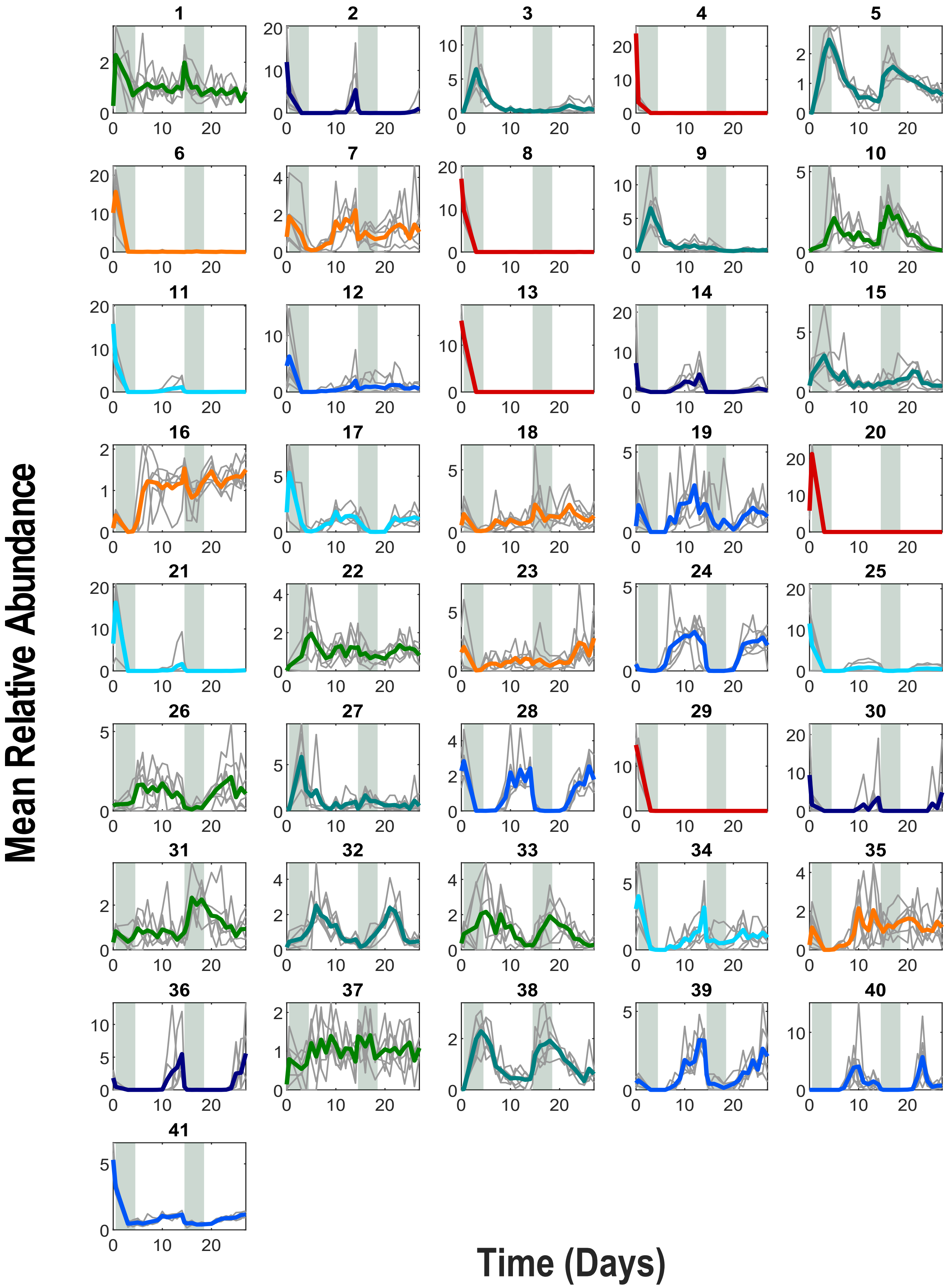
